# Inferring phylogenetic trees from mutational distance of transposable elements

**DOI:** 10.1101/846949

**Authors:** Juanjo Bermúdez

## Abstract

There are many methods for establishing a phylogeny available to researchers. Some of these are based on the mutational distance between ortholog sequences of DNA, and, from these, some are based on the analysis (presence/absence) of transposable elements in specific loci of ortholog sequences. We present a new approach: a method for inferring a phylogeny based on the mutational distance of transposable elements distributed along any segment of DNA. Our method doesn’t require previously having ortholog segments of DNA from every organism. The method can be fully automated, not requiring any previous or posterior data analysis nor data preparation.

## I. INTRODUCTION

Transposable elements (TEs) are repetitive DNA sequences that can amount to over two-thirds of the human genome [2][3] in some eukaryote species. These sequences become duplicated in the DNA under some circumstances by the action of polymerase proteins encoded in some of these TEs. Some scientists believe that this mechanism of replication has no evolutive purpose and are just an accident similar to viruses. Some others are starting to find their importance in some regulatory behaviors [8] [9] and as a force for evolution [10] (many existing genes were formed by the incrustation of TEs into other genes [4]).

SINEs (Short INterrpersed Elements) are a special type of TEs. SINEs are the only TEs that are non-autonomous by nature. They are small (80 - 600 base pairs) and rely on functional TEs for their replication. SINEs can be found in very diverse eukaryotes and have accumulated to impressive amount in mammals, where they represent between 5 and 15% of the genome with millions of copies.

SINEs have some common characteristics. They are relatively short. Most of these sequences are not subject to conservation given that they don’t form part of any coding or regulatory region: they are just copied at (usually) random locations in the DNA, forming part of the, so-called, junk DNA. Therefore, in most cases, mutations in SINE sequences are caused exclusively by replication errors and are not corrected by selection, becoming a quite reliable record for calculating elapsed time between replications. They also have the advantage of being quite more frequent replication events than vertical transmissions of genomes, therefore being a more abundant source of homolog sequences for the same historical period: you can compare multiple SINE sequences at different loci to find evolution patterns while genes are way more limited in number and subject to conservation. Given that SINEs are replicated intra-specimens at random loci, being originally exact copies of the original sequence, species derived from the species in which replications occurred will inherit multiple copies of the same sequence, inheriting, therefore, a multiple record of the same evolution event and not requiring the conservation of two concrete sequences in the same loci for comparison.

All these characteristics make SINEs a very interesting subject for phylogenetic studies but until now most efforts were only focused on distinguishing the presence or absence of a concrete SINE as a marker for insertion or deletion events.

We present a method for inferring phylogenetic trees based on the mutation distance between randomly selected TE elements (usually SINEs) distributed along different genomes.

## II. METHOD

### 1. Finding TE/SINE sequences

First, we make a search for potential TE/SINE sequences on every available DNA data. We make the search using consensus sequences for different target TE/SINE elements as query. For example, for mammals, we could use Alu, B1, Mare3, and Sine3. For birds, we could use Alu, B1, and Cr1e5 (not a SINE but an LTR TE). It works better if the TE/SINE sequences are known to be active in some of the species for which the tree will be applied as there will be more copies of the TE.

We have made use of the SLAST [6] tool for that step but in theory, any other local alignment search tool (like BLAST [5]) could have been used. The tool we used has the advantage that it already returns sequences with similar lengths, not discarding the subsequences that don’t match the query sequence. That’s convenient for the multiple alignment step and would have been needed to be done in an additional step in case of having used any other tool like BLAST.

### 2. Generating a multiple alignment of the sequences

Once we have the sequences, we need to align these for accurate comparison. Usually, the examples in this document will make use of around 200 sequences per species and TE/SINE class and around 10 species per study. That usually implies aligning between 1000 and 2000 sequences per TE/SINE class. But, e.g., the tree for the modern human populations (fig. 8) makes use of around 9.000 sequences per TE/SINE class; the overall study totaling around 30.000 sequences extracted from 36 individuals in the HGDP database [15].

**Fig. 1.**
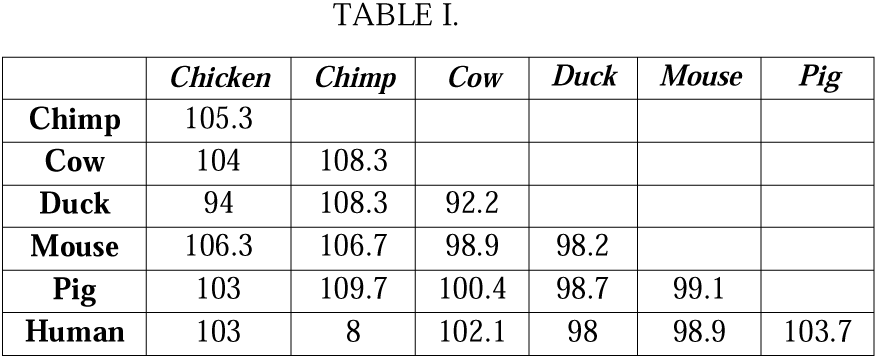
Mean of minimal distances for Mare3 SINEs.

**Fig. 2.**
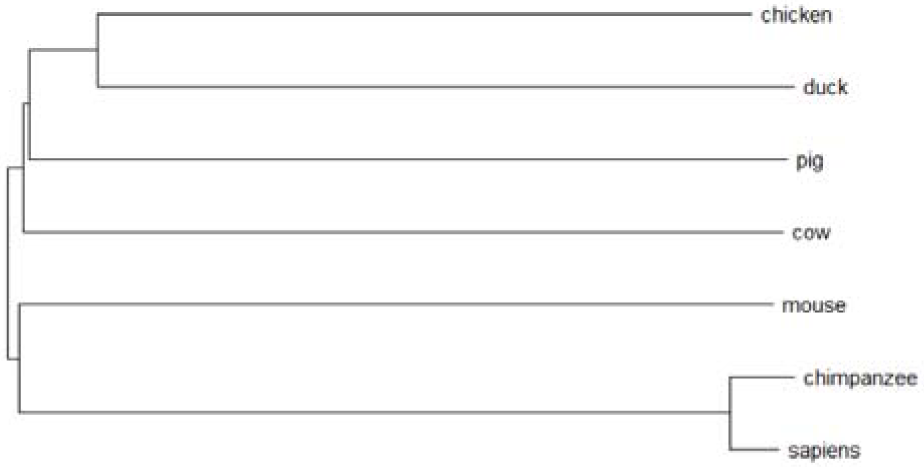
Phylogenetic tree generated from Mare3 element distances of Table I.

**Fig. 3.**
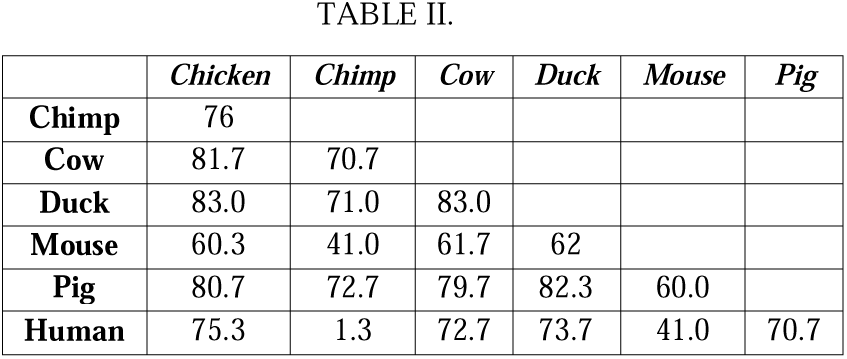
Minimal distances for B1 SINEs.

**Fig. 4.**
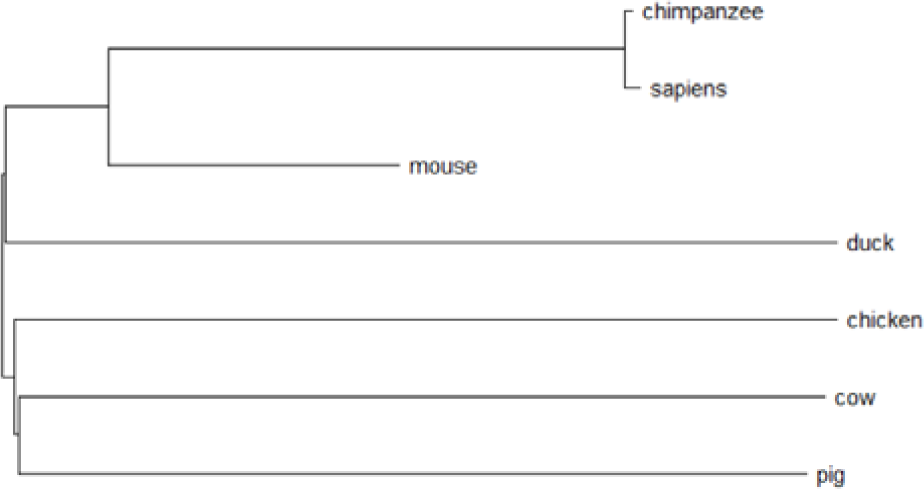
Phylogenetic tree generated from B1 element distances of Table II.

**Fig. 5.**
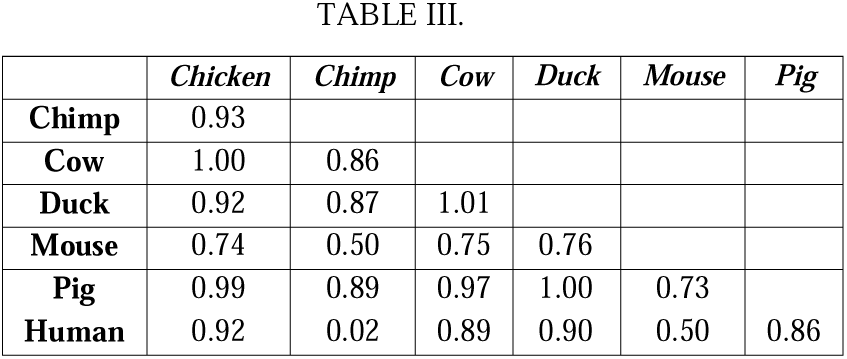
Table combining minimal distances for B1 and Mare3 SINEs.

**Fig. 6.**
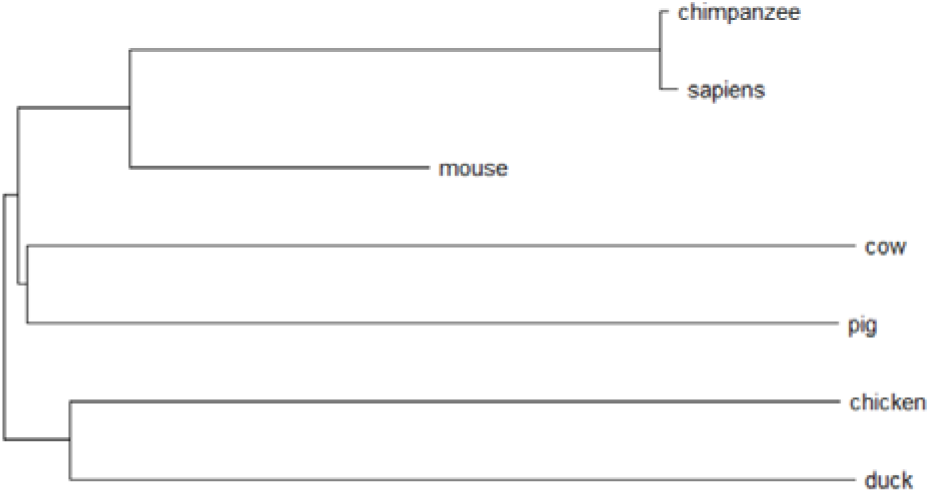
Phylogenetic tree generated from distances on Table III.

**Fig. 7.**
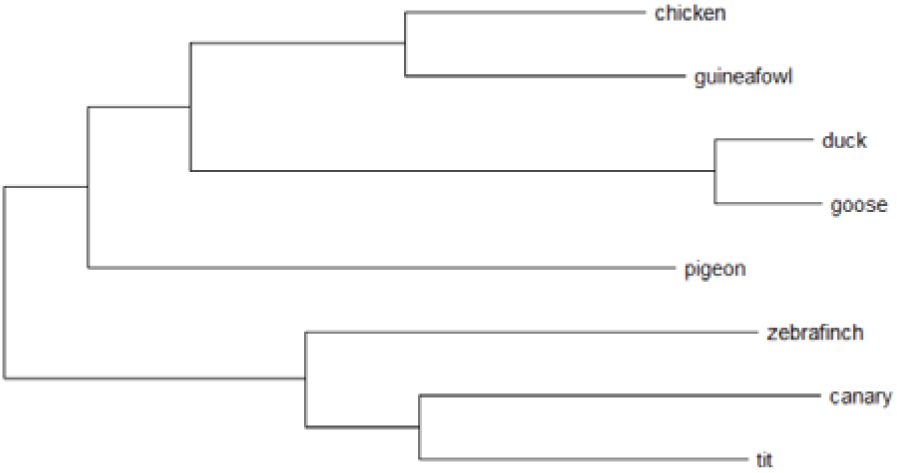
A tree obtained merging trees for Alu, B1, Cr1e5, Tgusine1, Sine3, Ther1, Mare3, Lf and 5ssaura TEs with data from the forward strand of different DNA fragments from each species.

**Fig. 8.**
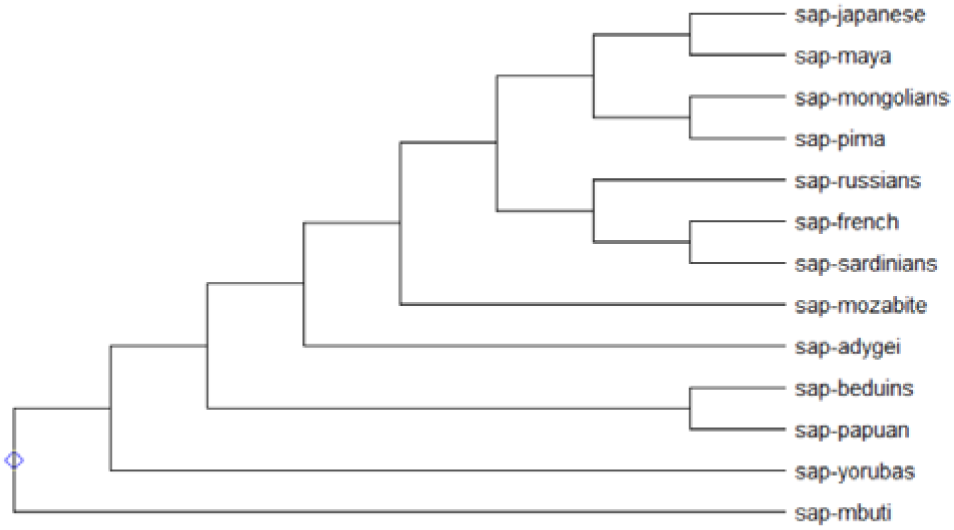
Tree (not representing real distances) obtained merging trees for Alu, Alusq10, and Cr1e5 with data from the first 50M bps of the forward strand of the chromosome 1 of 3 modern humans from each population in the tree.

### 3. Finding pairwise minimal distances

Once we have aligned the sequences, we compare pairwise distances from every sequence to every other sequence. We annotate the minimal distances from every species/organism to every other species/organism. In theory, the minimal distance between two TEs from different species indicates the number of mutations from the most recent ancestor of both species, but some circumstances can alter that distance. For instance, if the sequence is inside a conserved region the distance will be far lower (even zero). For that reason, instead of getting just the minimal distance we get an arbitrary number (n) of minimal distances and we calculate the mean. That helps to mitigate local effects.

With that information, we make a table. That table will be used for generating the phylogenetic tree. We can use algorithms like Neighbour Joining, Minimum Evolution or UPGMA for generating the trees from the distance table.

For instance, Table I shows the mean of minimal distances for Mare3 sequences between some vertebrates. We used sequences obtained from the forward strand of chromosomes 1 and 2 of each species.

Fig. 2 shows the corresponding tree for Table I.

### 4. Combining data from several trees

Next, we can combine information from different trees.

The minimal distance between two species for every

TE/SINE class should be caused by the same evolution split event but some circumstances could alter our results:

a. Some branches could have lost copies of some TEs due to deletion events.
b. There are no matching copies (or not enough) of the sequence in the segment of DNA that we used for the study.
c. We filtered the most significative matches for every organism in order to optimize resources but this way we selected different sets for every organism and some homologs were filtered out in some cases.
d. The TE remains active in one branch of the tree but not in other ones (i.e. ALU’s active in primates but not in other mammals). Consequently, we have a lot of short distances in that branch but not in other branches.

Therefore, if we take the minimal distance from many tables, we should get a better approximation to the real tree than using a single table given that we are avoiding possible circumstances that take to a higher distance in some of these tables. There is only one more consideration to have into account: the equivalent distances on every table don’t need to be the same, as the distance is proportional to the length of the TE sequence. But we can normalize distances dividing every distance by the maximal distance and these normalized distances should be almost proportional.

For instance, in Table II we have the mean of the minimum distances for the same species in Table 1 but making use of B1 as reference SINE sequence (also for forward strand of chromosomes 1 and 2).

Combining normalized minimal distances from Table I and Table II we get Table III.

## III. RESULTS

Some phylogenetic trees obtained with this method are shown in figures 7, 8 and 9.

**Fig. 9.**
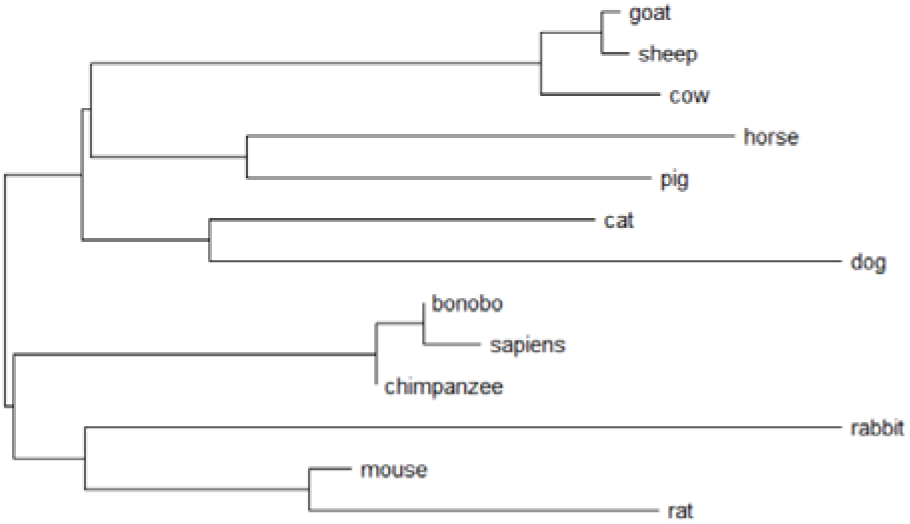
Tree obtained merging trees for Alu, B1, Can, Mare3, Sine3, Id and Bovta SINEs with data from forward strand of chromosomes 1, 2, 3, 4, 5 and 6.

As we can see, the trees are quite good approximations to the actually established phylogenies for these species. It even approximates fairly well the phylogeny of old ancestors of the species: it classifies, for example, rodents related to the rabbit and near to primates.

Everything that was needed for generating these trees was a fragment of the genome of the species/organisms in FASTA format. No data preparation or human intervention was needed apart from selecting the TEs for the study (which would not be necessary for adding new species to the studies).

## IV. OBSERVATIONS

Apart from the results of this work, some other observations are noticeable for further studies.

### A. Ancestral origin

Even if some SINE’s are specific for some species, far more distant species usually retain also a proportional distance relative to their phylogeny relation. That seems to indicate that even if some SINEs are activated at some point by a specific species, they are not created from scratch but as a derivation of any very ancient SINE and that’s why other species not derived from that reactivation event also retain a proportional distance. In these cases, the phylogenetic structure is retained but there is a scaling effect on the tree making the tree more compressed in the regions for species originated at events nearer to the reactivation event and making it more dilated for the most distant species.

We can see in fig. 10, for example, how goat and sheep seem to be in appearance more distant from a common ancestor than they really are, while sapiens, chimpanzee and bonobo seem to be far closer. But even that, their phylogenetic relationships with other nodes and their location in the tree is preserved quite well. Considering that AluSq10 is specific for primates, that seems to indicate that even species not derived from the common ancestor of primates have remains of copies derived from an ancestor of AluSq10. That seems to indicate that the common ancestor for these sequences is older than the common ancestor of the species in the tree. Methods for normalizing that scale effect (apart from merging trees) should be studied in the future.

**Fig. 10.**
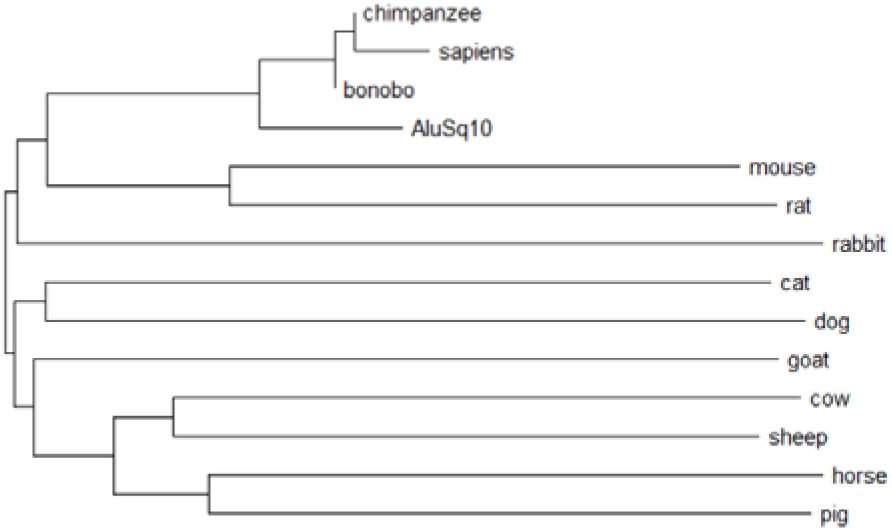
Mammals tree obtained from AluSq10 mutation distances with data from the forward strand of chromosomes 1 and 2.

### B. Multiple Alignment

Multiple Alignment (MA) is an important step for our method as it is what determines the quality of the comparisons between sequences. Using pairwise alignments would also be a valid approach (even better) but was discarded in this study due to the reduced speed performance that implies.

We have made tests with two Multiple Alignment algorithms: Clustal Omega [11] and a relatively simple algorithm we specifically designed for non-conserved sequences.

We observe that:

1. Results are slightly different depending on the MA algorithm in use.
2. Clustal Omega seems to retain relations better between close species but the other one seems to retain relations better for more distant species.
3. Overall, both algorithms have similar results when enough data is used for the study, having not found a significative difference in the amount of data needed for each algorithm. But if one has to be classified as having a better performance it would be the one we designed specifically. All trees in this document were produced making use of our specifically designed MA algorithm.

### C. Classification of species based on partial DNA samples

This method, in theory, makes possible to produce an automated tool for classifying any species for which we only have a fragment of DNA. We would insert that sequence and obtain as result a phylogenetic tree including the location in the phylogeny of the species corresponding to the DNA fragment. Further studies will be implemented to test the requirements and capacities for such a tool.

### D. Distances and “dancing” branches

Further analyses of the reliability of the distances of the nodes in the trees are needed. We can observe in fig. 9 how the ancestor of primates and rodents is quite near to the common split with the rest of mammals. A slight alteration of the distances could take to a different tree in which rodents and primates would not be in the same branch. That is in line with some controversies about the exact location of primates in the mammals’ tree and could indicate that the proportion of distances generated by this method could be reliable.

## V. CONCLUSIONS

We demonstrate that it’s possible to infer phylogeny from the analysis of mutational distances between TE elements in different species or organisms. The generated phylogenies, indeed, are in concordance with other studies.

Our method has some advantages:

- It is completely automated: there is no need for data preparation or analysis. Only a fragment of a DNA sequence is needed.
- It’s possible to establish phylogeny starting from any segment of DNA from each species: no need to have aligned orthologs from every species.
- We make use of data from multiple loci, diminishing this way possible local repression effects at some loci, which are usually a concern using other methodologies.

This method could also be derived into a tool to place samples of DNA of uncertain origin into a phylogeny.

## Supporting information

Some extracted data, alignments and trees.

## VI. ACKNOWLEDGMENTS

The authors of this research adhere to the Agile Science Manifesto v 1.0 [7] and this document was written in concordance with its arguments.

Some data for this study were obtained from [13][14][15][16][17][18].

Some free software tools used were: [19][20][21]

